## Supplemental Material for "Computationally designed GPCR quaternary structures bias signaling pathway activation"

**Supplementary Table S1. Distribution of closed and open CXCR4 dimer conformations.** The dimer interface energy in Rosetta Energy Units are reported for the best open-dimer or closed-dimer conformations adopted by each CXCR4 variant modeled in the inactive or active state.

| | INACTIVE STATE | | ACTIVE STATE | |
|---|---|---|---|---|
| | Dimer Interface Energy | | Dimer Interface Energy | |
| | Open-dimer | Closed-dimer | Open-dimer | Closed-dimer |
| WT | -24.28 | -18.38 | -25.39 | -21.06 |
| N192$_{ECL2}$W | -22.67 | -22.16 | -19.52 | -25.09 |
| W195$_{5.34}$L | -23.59 | -21.53 | -21.3 | -26.07 |
| L194$_{5.33}$R | -26.68 | -20.83 | -24.7 | -20.75 |

**Supplementary Table S2. Relationship between calculated dimerization propensity and dimerization BRET signals measured for CXCR4 variants.** The predicted dimerization propensities of the CXCR4 variants relative to the WT receptor are calculated from the energies of the dimer conformations in the inactive and active states (**see Methods**). The net dimerization BRET signals (**see Fig.2**) normalized to WT are provided for comparison.

| | INACTIVE STATE | | ACTIVE STATE | |
|---|---|---|---|---|
| | Predicted dimerization propensity | Constitutive net BRET | Predicted dimerization propensity | Agonist induced net BRET |
| WT | 1.00 | 1.00 | 1.00 | 1.00 |
| N192$_{ECL2}$W | 0.42 | 0.57 | 0.76 | 0.69 |
| W195$_{5.34}$L | 0.65 | 0.81 | 1.76 | 1.22 |
| L194$_{5.33}$R | 7.57 | 0.95 | 0.56 | 0.88 |

**Supplementary Table S3. Distribution of open and wide-open μOR dimer conformations.** The dimer interface energy in Rosetta Energy Units are reported for the best open dimer or wide-open dimer conformations adopted by each μOR variant modeled in the active state.

| | ACTIVE STATE | | | | | |
|---|---|---|---|---|---|---|
| | Dimer Interface Energy | | Beta arrestin binding energy | | Gi binding energy | |
| | Open-dimer | Wide-open dimer | Open-dimer | Wide-open dimer | Open-dimer | Wide-open dimer |
| WT | -27.9 | -22.7 | -9 | -6.3 | -10.9 | -9.7 |
| W230$_{5.34}$A | -26.7 | -23.3 | ND | ND | ND | ND |

**Supplementary Figure S1. Workflow of the computational approach.** Starting from homolog templates, ligand-bound GPCR inactive and active state monomers are generated using iPHoLD[31]. Monomers are assembled into GPCR dimers and into complex with G-protein or β-arrestin by flexible docking. Loop and residue motifs are designed at the predicted dimer interface to destabilize (in this example) or stabilize selectively a specific dimer conformation (e.g. open-dimer). Designed monomers are assembled into GPCR dimers and into complex with G-proteins or β-arrestin to assess the shift in dimer conformations distribution (calculated as the energy difference between the closed and open designed structures, $\Delta E^{(c\text{-}o)}{}_i$) and the associated functional (i.e. G-protein versus β-arrestin binding) shift. The design-quaternary structure assembly cycle is repeated until a significant shift is achieved, i.e. $\Delta E^{(c\text{-}o)}{}_i$ and $\Delta E^{(c\text{-}o)}{}_a$ << 0 providing monomer stability is not affected significantly.

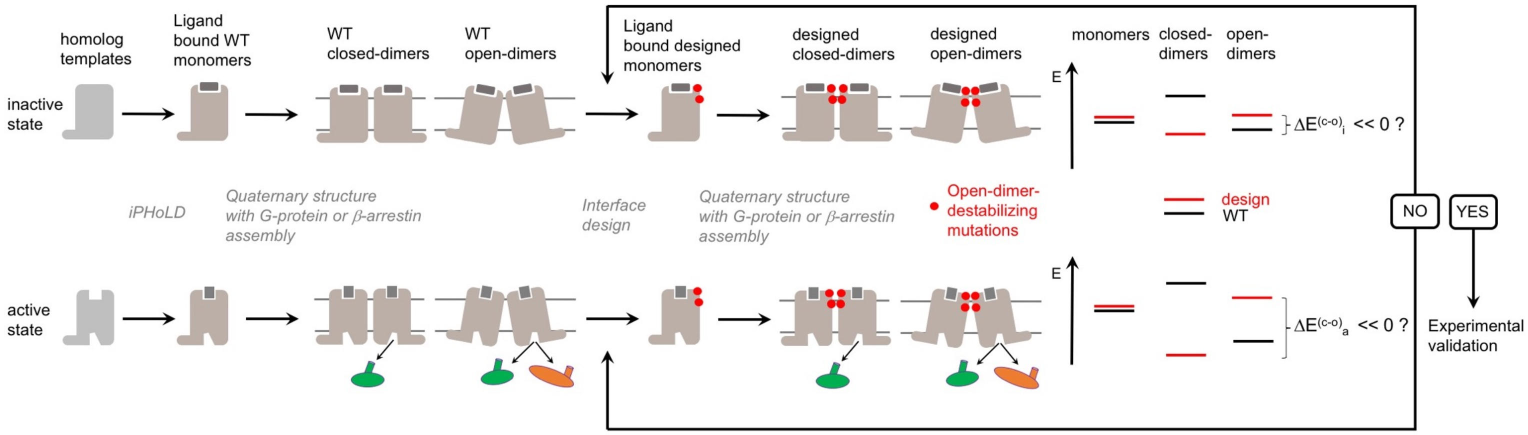

**Supplementary Figure S2. Geometrical comparison between the open and closed CXCR4 dimer conformations. Top.** Superposition between closed and open dimer conformations (the aligned monomer of the closed form is omitted for clarity). TM helices IV and V are labelled in each protomer. The Cα rmsd between the open-dimer inactive state model and the antagonist bound X-ray structure (3odu) is 3.5 Å over the entire dimer structure. **Bottom.** The interhelical angle and distance between the TM helix 5 of each monomer are used to characterize the dimer conformation. A vector is fitted to the Cα coordinates of each TM helix 5. The angle and mid-point distance between the two vectors are measured. Two representative models of the open-dimer (θ = 53° and d = 13.6Å, left) and closed-dimer conformation (θ = -8.2° and d = 21.4Å, right) are shown.

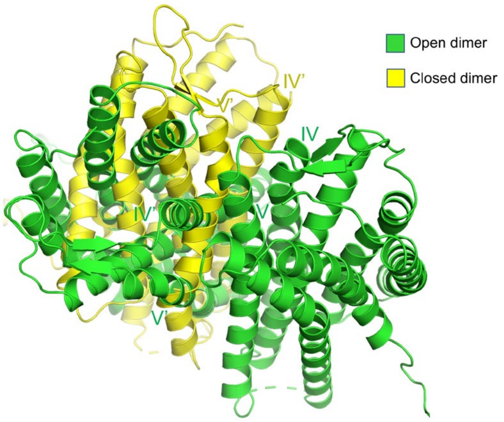


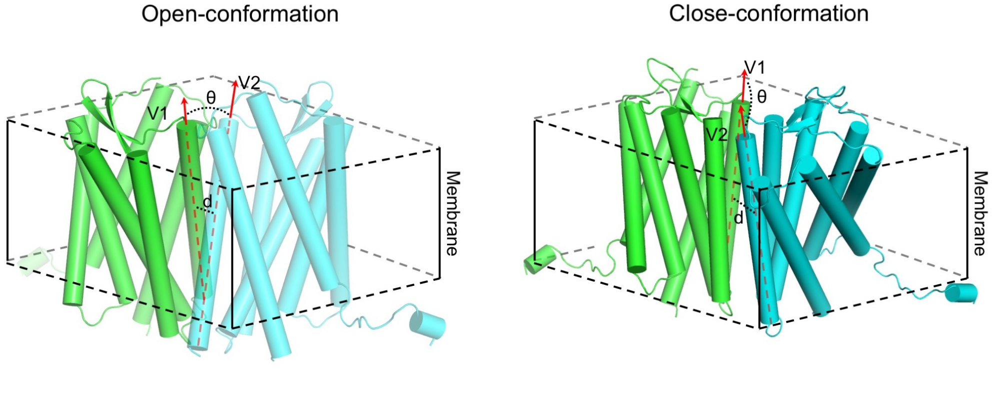


| | INACTIVE STATE | | | | ACTIVE STATE | | | |
|---|---|---|---|---|---|---|---|---|
| | Angle (deg) | | Distance (Angstrom) | | Angle (deg) | | Distance (Angstrom) | |
| | Open-dimer | Closed-dimer | Open-dimer | Closed-dimer | Open-dimer | Closed-dimer | Open-dimer | Closed-dimer |
| WT | 60.62 | -8.86 | 14.24 | 19.71 | 51.54 | 26.59 | 14.45 | 20.62 |
| N192W | 55.07 | -10.90 | 13.92 | 22.09 | 59.74 | 29.14 | 12.61 | 21.57 |
| W195L | 74.18 | 1.35 | 13.10 | 21.83 | 55.82 | 35.03 | 12.27 | 21.43 |
| L194R | 58.49 | -17.45 | 15.46 | 23.66 | 50.35 | 23.72 | 12.72 | 21.44 |

**Supplementary Figure S3. CXCR4 strongly interacts with β-arrestin in the open-dimer conformation only.** Representative lowest energy open and closed dimer conformations of CXCR4 WT in the active state bound to β-arrestin (a) and Gi (b). The structures with the lowest binding energy between the receptor and each effector are shown with a zoomed view of the main binding interface. The binding energies between the CXCR4 conformations and β-arrestin or Gi are provided in Rosetta Energy Units (REU). The closed-dimer conformation prevents optimal binding of the β-arrestin's finger loop with the CXCR4 monomer intracellular binding groove. TM helices are labelled on each CXCR4 protomer.

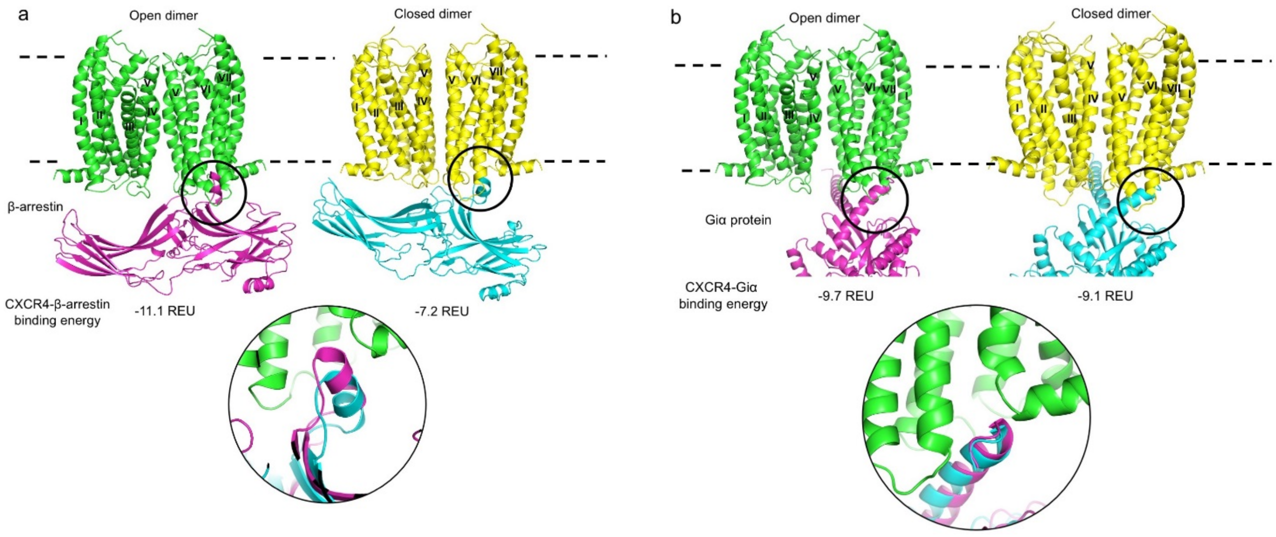

**Supplementary Figure S4. Steric hindrance prevents CXCR4 in the closed dimer conformation to strongly interact with β-arrestin. a.** The closed-dimer conformation is aligned to the β–arrestin-bound open-dimer receptor complex by superimposing one CXCR4 monomer. Open- and closed-dimer conformations are colored in green and yellow, respectively. The docked β–arrestin is colored in magenta. The two insets highlight regions of close contacts between the β–arrestin and the open-dimer CXCR4 (**b:** Helix 8 of CXCR4 monomer 2 with the C-tip of β-arrestin**, d:** ICL2 of CXCR4 monomer 1 with the C-loop of β-arrestin) that would be disrupted in the closed-dimer conformation because of major steric clashes between the β–arrestin and the receptor second monomer (**c, e**). TM helices are labelled on each CXCR4 protomer.

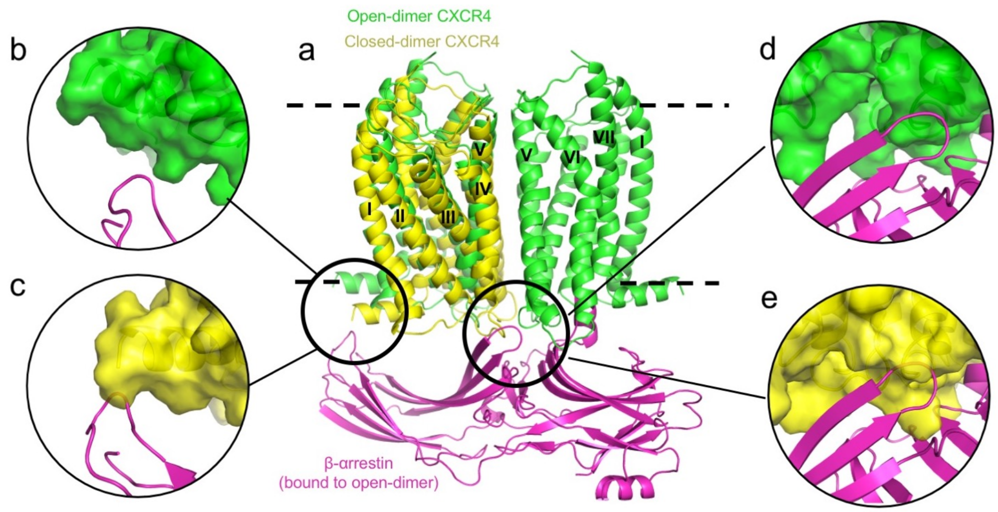

**Supplementary Figure S5. Correlation between closed dimer stabilization and BRET signal changes upon receptor activation.**

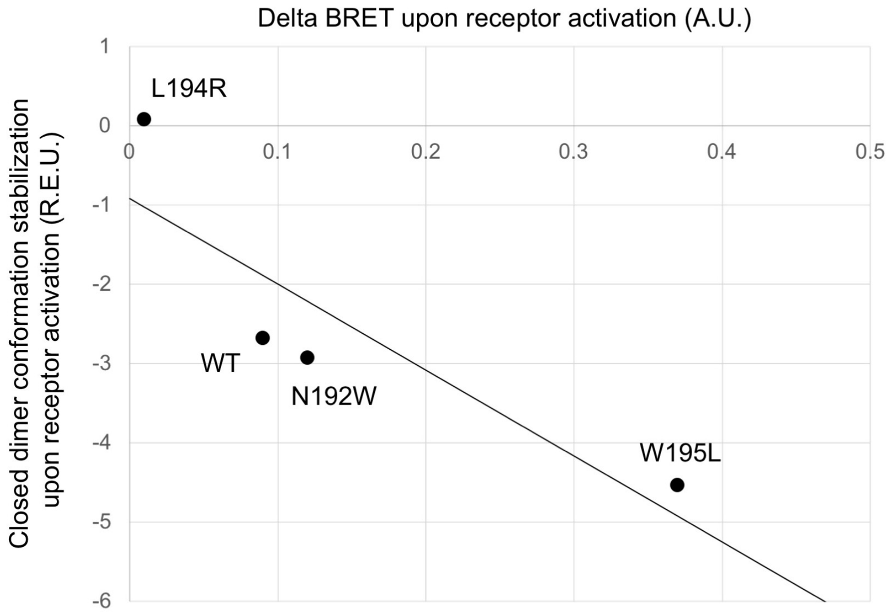

**Supplementary Figure S6. Cell surface expression of CXCR4 variants. a-c.** Cell surface expression of Myc-CXCR4-RLuc (**a**), HA-CXCR4-YFP (**b**) and HA-CXCR4 (**c**), as well as their respective mutant forms, was determined by ELISA using an anti-Myc (**a**) or anti-HA (**b, c**) antibody. **d.** Cell surface expression of endogenous WT CXCR4 in U87 versus HEK293-T cells was determined by Flow cytometry using an anti-CXCR4 antibody. **e.** Cell surface expression in U87 cells of exogenous HA-CXCR4, WT or W195$^{5.34}$L, was determined by Flow cytometry using an anti-CXCR4 (left) and HA antibody (right).

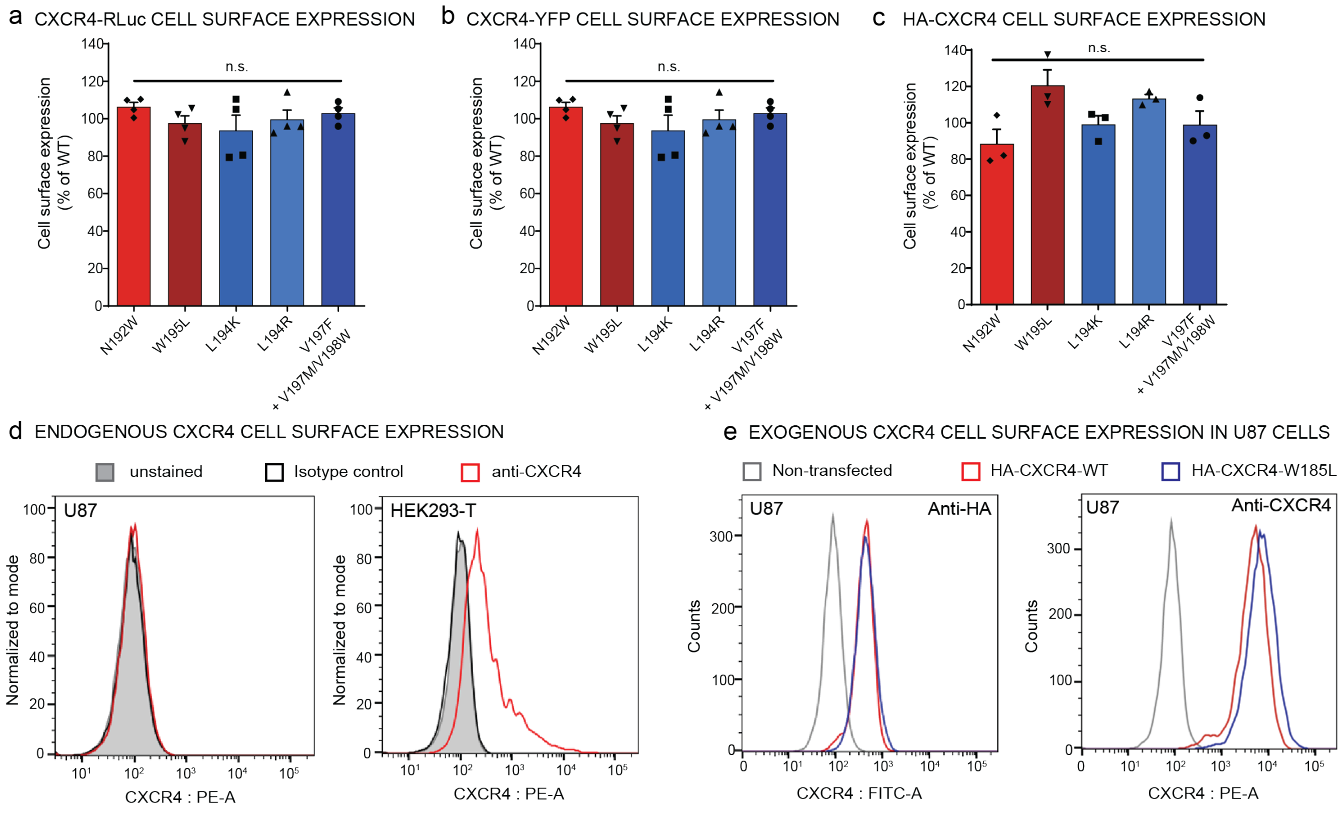

**Supplementary Figure S7. Titration curves of CXCR4 association**. CXCR4 association BRET titration curves were performed in cells co-transfected with a constant amount of CXCR4-Rluc and increasing amounts of WT or mutant forms of CXCR4-YFP, as indicated. $BRET_{480\text{-}YFP}$ was measured after the addition of coel-h (10 min) and 200 nM CXCL12 or vehicle (15 min). The curves shown are derived from individual titration curves that are representative of three independent experiments. The error bars represent SD from triplicate wells.

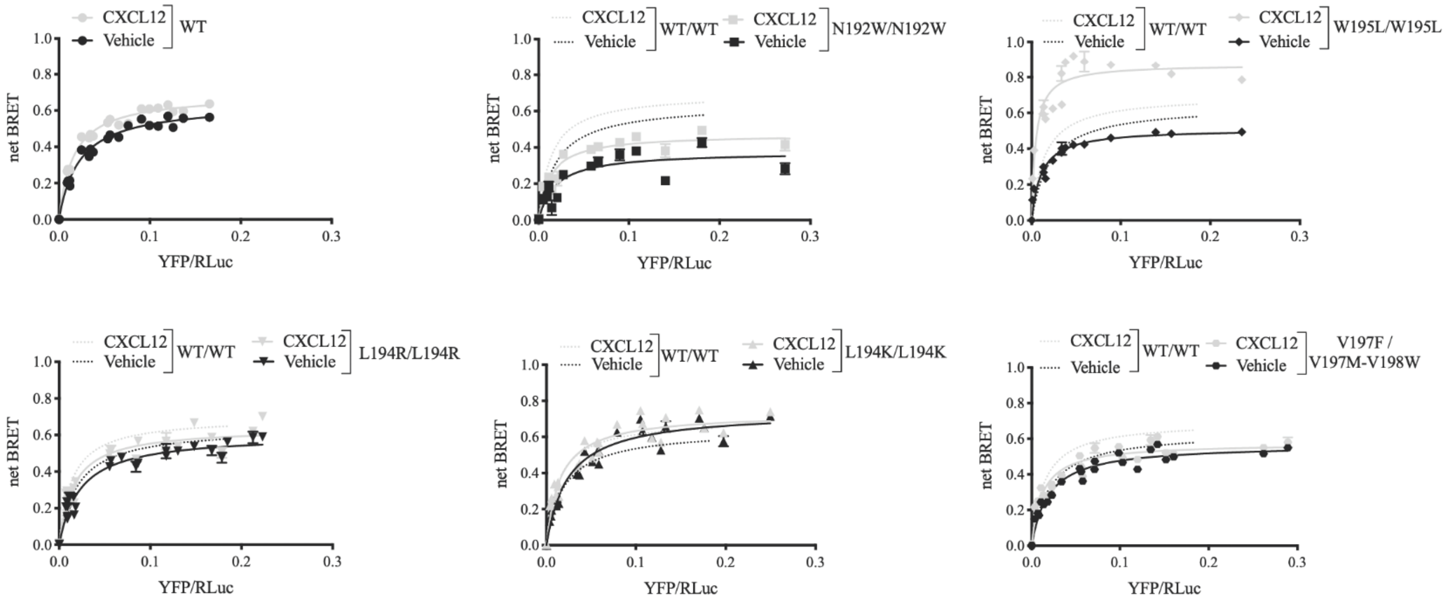

**Supplementary Figure S8. Translocation of β-arrestin2 to the plasma membrane**

(Left) Schematic representation of the ebBRET-based assay used to follow agonist-induced βarr2 recruitment to the receptor by monitoring the interaction between βarr2-RLucII and rGFP-CAAX. (Right) Cell surface translocation of βarr2 was assessed by BRET400-GFP10 in HEK293T transfected with HA-CXCR4, WT or mutant as indicated, βarr2-RLucII and rGFP-CAAX. BRET400-rGFP between βarr2-RLucII and CAAX-rGFP was measured after the addition of coel-400a (5 min) and CXCL12 (15 min). Data are expressed as agonist-promoted BRET (ΔBRET). CXCR4 mutations predicted to stabilize the open-dimer or the closed-dimer conformation are annotated with a blue or red dimer symbol, respectively. Data shown represent the mean ± SEM of at least three independent experiments.

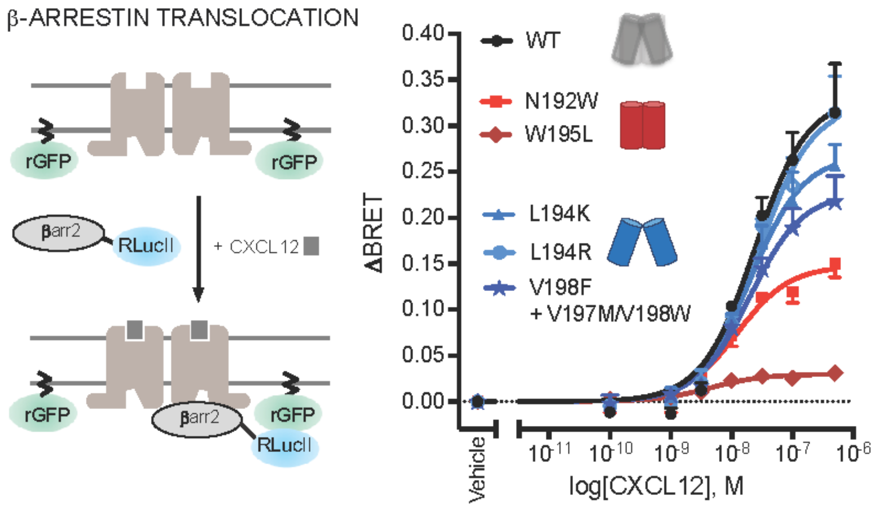

**Supplementary Figure S9. Geometrical comparison between the open and closed μOR dimer conformations. Top.** Superposition between the Open and Wide-open dimer conformations (the aligned monomer of the Wide-open form is omitted for clarity). TM helices V and IV or VI are labelled in each protomer. **Bottom.** The interhelical angle and distance between the TM helix 5 of each monomer are used to characterize the dimer conformation. A vector is fitted to the Cα coordinates of each TM helix 5. The angle and mid-point distance between the two vectors are measured. Two representative μOR WT models of the Wide-open-dimer (θ = 45° and d = 13.7Å, left) and Open-dimer conformation (θ = 30° and d = 11.8Å, right) are shown.

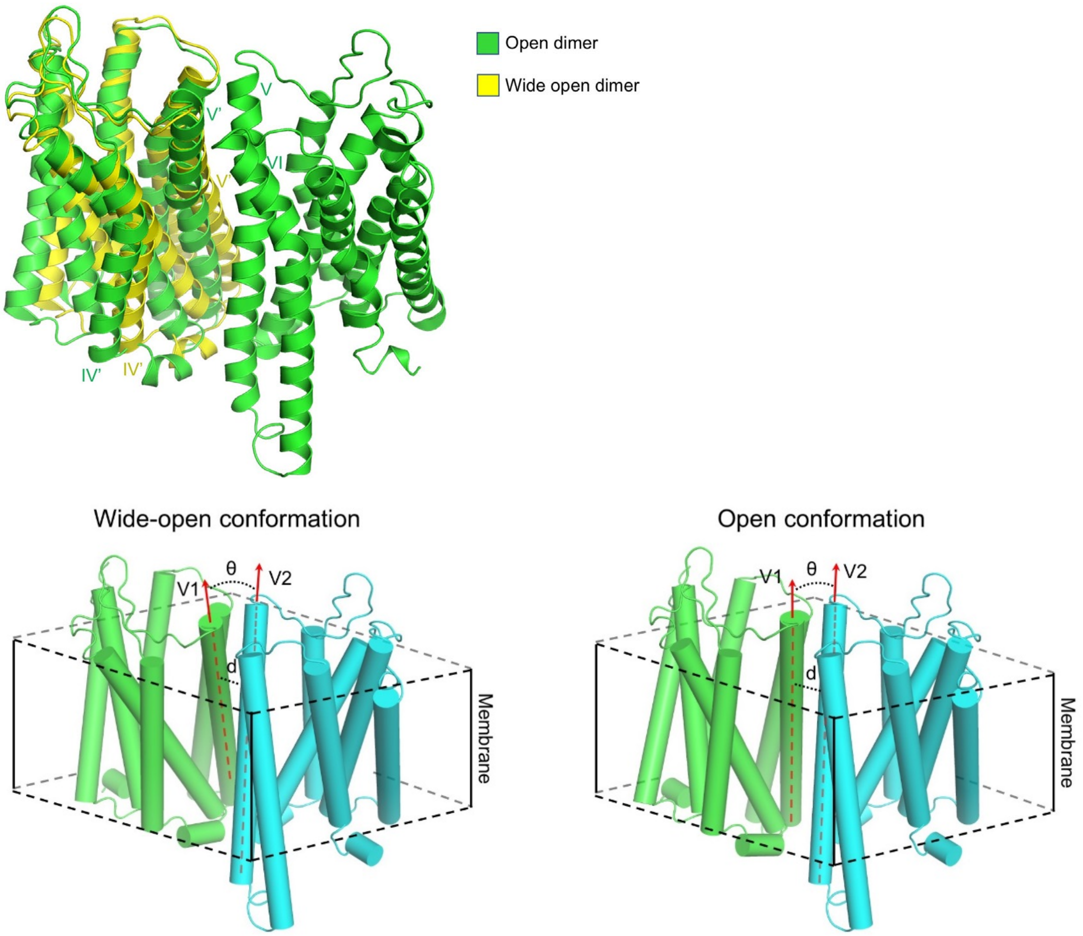


| | ACTIVE STATE | | | |
|---|---|---|---|---|
| | Angle (deg) | | Distance (Angstrom) | |
| | Open-dimer | Wide open-dimer | Open-dimer | Wide open-dimer |
| WT | 30.42 | 44.62 | 11.80 | 13.70 |
| W5.34A | 29.75 | 43.79 | 11.70 | 13.86 |
